# Release from cell cycle arrest with Cdk4/6 inhibitors generates highly synchronised cell cycle progression in human cell culture

**DOI:** 10.1101/2020.07.04.187625

**Authors:** Eleanor Wendy Trotter, Iain Michael Hagan

## Abstract

Each approach used to synchronise cell cycle progression of human cell lines presents a unique set of challenges. Induction synchrony with agents that transiently block progression through key cell cycle stages are popular, but change stoichiometries of cell cycle regulators, invoke compensatory changes in growth rate and, for DNA replication inhibitors, damage DNA. The production, replacement, or manipulation of a target molecule must be exceptionally rapid if the interpretation of phenotypes in the cycle under study are to remain independent of impacts upon progression through the preceding cycle. We show how these challenges are avoided by exploiting the ability of the Cdk4/6 inhibitors, palbociclib, ribociclib and abemaciclib to arrest cell cycle progression at the natural control point for cell cycle commitment: the restriction point. After previous work found no change in the coupling of growth and division during recovery from CDK4/6 inhibition, we find high degrees of synchrony in cell cycle progression. Although we validate CDK4/6 induction synchronisation with hTERT-RPE-1, THP1 and H1299, it is effective in other lines and avoids the DNA damage that accompanies synchronisation by thymidine block/release. Competence to return to cycle after 72 hours arrest enables out of cycle target induction/manipulation, without impacting upon preceding cycles.

## Background

Synchronised progression through the cell division cycle throughout a population supports the ability to extrapolate the biochemical and functional attributes of the synchronised bulk population back to infer behaviour in an individual cell [1,2]. Many approaches are popular. Bulk levels of DNA or cell cycle markers support fractionation of live, or fixed, cell populations into pools enriched for discrete cell cycle stages [3,4]. Although yields are low, selection synchronisation based upon size, or mitotic shake off, are highly effective approaches to isolate cells in one cycle phase from a large population of asynchronous cells [5,6]. However, the ease of induction synchrony makes it the most widely applied approach.

Induction synchrony exploits the ability of transient exposure to a particular context to accumulate cells at a discrete cell cycle stage, before removal of the context simultaneously releases all cells in the population, to progress synchronously through subsequent phases of the cell division cycle [1]. In yeasts transient ablation of cell cycle regulators through reversible conditional mutations and the addition of mating pheromones predominate [7,8]. Although the advent of analogue sensitive versions of cell cycle kinases has introduced analogous chemical genetic approaches into human tissue culture studies [9–12], induction synchrony via serum starvation [13], or activation of either the DNA replication or spindle assembly checkpoints [5,14] remain the most widely used.

The discovery that transient treatment with thymidine synchronised mitotic progression [14], led to protocols that sharpened the degree of synchrony by imposing a second thymidine block before releasing cells into the cycle of study [15–18]. This “double thymidine block” remains one of the most popular choices, however, the power of this approach, its reliance upon the DNA replication checkpoint to arrest S phase progression with stalled DNA replication forks, comes at a cost. Although the cell cycle arrest is robust in many lines, the stalled forks are prone to collapse over the extended arrest and subsequent attempts at repair introduce damage and chromosomal rearrangements [19–21]. There are also reports of understandable impacts upon RNA biology during the long block [22,23]. Thus, this popular approach can be of limited utility in the study of S phase, some transcriptional and chromatin associated events.

When early cell cycle events are to be analysed, induction synchrony via release from a mitotic arrest in the previous cell cycle, provides an attractive alternative. However, like other forms of induction synchrony that rely upon an arrest within the cell division cycle, prolonged cell cycle arrest will generate an imbalance in the many regulators, whose levels fluctuate with cell cycle progression as a consequence of stage dependent transcription and/or destruction [24]. Consequently, the next cycle may well be altered by excessive regulatory activities, or substrates, inherited from the preceding, arrested, cycle. Incisive studies by Ginzberg and colleagues revealed how counter-measures to accommodate some imbalances promote adjustments in growth rates at two points in the cycle [25]. Prolonged mitotic arrest can also initiate apoptotic pathways [26] and/or leave a memory of the mitotic arrest that modifies cell cycle progression in the next cycle and beyond [27–30].

Thus, while highly informative for some questions, data obtained through traditional induction synchrony approaches, that rely upon arrest *within* the cycle, have to be interpreted with caution. They must be consolidated with complementary data from alternative approaches, to reveal the commonalities that exclude the artefacts incurred in one given approach to synchronisation.

A further challenge in synchronising cell cycle progression throughout a population arises when there is a need to assess the impact of protein depletion, induction or replacement. It is critical to ensure that the destruction, induction, or activation of a mutant variant starts after the synchronising procedure is complete. If not, then the phenotype can be a legacy arising from perturbation of progression through the previous cycle, rather than a direct impact upon the cycle being studied. Advances in degron and PROTAC technologies may overcome many of these challenges [31–33]. However, even with many induction synchronisation approaches, the switch from one version of a protein to another must be exceptionally rapid and complete if perturbation of the preceding cycle is to be avoided.

Inspired by the power of pheromone induction synchronisation at G1 phase of yeast cell cycles [8,34], we explored the utility of induction synchrony with CDK4/6 inhibitors palbociclib, ribociclib and abemaciclib. These inhibitors arrest cell cycle progression of mammalian tissue culture cells at the restriction point in G1 phase prior to commitment to the cell cycle. Synchronisation by induction from the natural pause point in the cycle appeals because the cell cycle programme is yet to be set in motion and extended arrest via Cdk4/6 inhibition does not invoke compensatory changes in cell cycle or growth controls; rather it simply adjusts cell size control [25].

Cdk4 and Cdk6 kinases determine commitment to the cell cycle of many cells. They partner Cyclin D and the Kip family members p21 and p27 to generate active trimeric kinase complexes that phosphorylate the C terminus of the retinoblastoma (Rb) family proteins [35–41]. This mono-phosphorylation supports further phosphorylation of Rb by Cdk1/Cdk2-Cyclin E and Cdk1/Cdk2-Cyclin A complexes [10]. Hypo-phosphorylated Rb binds tightly to the transcription factors of the E2F family, to block the transcription of genes required for cell cycle commitment. Rb hyper-phosphorylation relieves this inhibition, to promote transcription of cell cycle genes, including Cyclins E and A. Induction of these Cyclins rapidly boosts Rb phosphorylation by Cyclin E and Cyclin A Cdk complexes [42]. Cyclin E and Cyclin A complexes seal commitment to the cycle by inhibiting the anaphase promoting complex/cyclosome (APC/C) activating component Cdh1, thereby stabilising APC/C^Cdh1^ targets, including Cyclin A [43–47].

The key role played by Cdk4-Cyclin D and Cdk6-Cyclin D in presiding over cellular proliferation, and the contrasting ability of mice to survive genetic ablation of Cdk4, Cdk6 and Cyclin D and the addiction of cancer lines to these kinases, prompted the development of clinically successful Cdk4/6 inhibitors [48–52]. These inhibitors bind to the inactive Cdk4-Cyclin D and Cdk6-Cyclin D dimers, rather than the active trimeric complexes, yet they impose a very efficient cell cycle arrest [41]. It is thought that this counter-intuitive impact arises from the sequestration of the inactive dimers, or monomeric kinases by the drugs. This sequestration releases p21 and p27 to elevate concentrations of these Cdk2 inhibitors, to a level where they block the ability of the Cdk2-Cyclin A and Cdk2-Cyclin E complexes to promote the feedback loops and S phase events that drive commitment to the cycle [41,53]. Each drug has a specific off target profile. Ribociclib’s impressive specificity is almost matched by palbociclib, while abemaciclib shows significant off-target impacts, with notable activities towards Cdk1, Cdk2, Cdk7 and Cdk9 complexes, that can induce an arrest in G2 alongside G1 [54]. Paradoxically, this off-target impact may account for abemaciclib’s greater efficacy in some clinical settings [54,55].

We describe how CDK4/6 induction synchrony generates highly synchronous progression through the cell cycles of a number of lines, without the marked appearance of a marker of DNA damage, foci of staining of phospho-γ-H2AX, that accompanies thymidine induction synchronisation. The ability to return to cycle after 72 hours arrest in G1 with palbociclib will support a broad range of manipulations in the arrested, non-cycling state [56]. Thus, any impacts upon progression through the cycle of study will not be a secondary consequence of perturbation of the preceding cell cycle.

## Methods

### Cell lines and Cell Culture

The following lines were purchased from ATCC at the start of the study: hTERT-RPE1 (ATCC Cat# CRL-4000, RRID:CVCL_4388), A-549 (NCI-DTP Cat# A549, RRID:CVCL_0023), THP-1 (CLS Cat# 300356/p804_THP-1, RRID:CVCL_0006) and NCI-H1299 cell line (NCI-DTP Cat# NCI-H1299, RRID:CVCL_0060). Upon receipt, lines were expanded and frozen into aliquots of 1 × 10^7^ cells that were expanded by at least two cycles of continual passage prior to each experiment. Unless otherwise stated, hTERT-RPE-1 and A549 were maintained in high glucose Dulbecco’s modified eagles medium (DMEM: D6546, Sigma Aldrich) supplemented with 10% FBS (Hyclone, South American Origin, SV30160.03, GE Healthcare) 2 mM GlutaMAX (Gibco, 35050061) and 1000 U/ml Penicillin/Streptomycin (Gibco, 15140122). THP1 and H1299 were maintained in Roswell Park Memorial Institute 1640 medium (RPMI) supplemented with 2 mM GlutaMAX (61870044, Gibco), 10% FBS (Hyclone, South American Origin, SV30160.03, GE Healthcare) and 1000 U/ml Penicillin/Streptomycin (Gibco, 15140122). Cells were maintained at a confluency of less than 70% and populations were not passaged more than 10 times before any experiment. In Figure 2e, where hTERT-RPE-1 cells are grown in RPMI, cells were originally grown from the frozen stock in DMEM, passaged twice in RPMI, then plated in RPMI at the start of the experiment.

### Drug treatment

Palbociclib (PD-0332991), Abemaciclib (LY2835219) and Ribociclib (LEE011) were purchased from Selleck (catalogue numbers S1116, S5716, S7440 respectively), while thymidine (T1895) and nocodazole (M1404) were from Sigma-Aldrich. Palbociclib, Abemaciclib, Ribociclib and Nocodazole were dissolved in DMSO to generate stock solution of 10 mM in each case. Thymidine was dissolved in water to make a stock solution of 100 mM. All stocks were stored in aliquots at −20 °C.

For adherent cell lines (RPE1, A549 and H1299), cells were released from the substrate by treatment with trypsin (15400054, Gibco), pelleted by centrifugation at 300 g for 5 minutes before resuspension in growth media. Cells were counted using a TC20 automated cell counter (BioRad) and 2.5 × 10^5^ cells were plated in a 10 cm dish (353003, Falcon) with 10 ml media (plating density = 4.4 × 10^3^ per cm^2^). For the THP1 suspension line, cells were pelleted by centrifugation at 300 g for 5 minutes, resuspended in growth media and counted, before 1 × 10^6^ cells were seeded in 10 ml media in a 10 cm dish. Cells were then incubated for 6 or 12 hours (see figure legends) before drug was added.

### Flow Cytometry

For cell cycle analysis, cells were washed once with phosphate buffered saline (PBS) before releasing from the substrate with trypsin, and washed in phosphate buffered saline before fixation in ice-cold 70% Ethanol and freezing at −20°C for at least 18 hours up to a maximum of 2 weeks. Fixed cells were pelleted, washed 3 times in PBS at room temperature before 50 μl 100 μg/ml RNase (NB-03-0161, Generon) was added to the final pellet, followed by 500 μl 50 μg/ml propidium iodide (P4170, Sigma-Aldrich) dissolved in PBS. Samples were analysed within a window of between 30 minutes and 6 hours after addition of propidium iodide. Data were acquired on a BD LSR II flow cytometer (BD Biosciences) using FACSDiva™ software (BD Biosciences) and analysed with FlowJo software (BD Biosciences). 1 × 10^4^ cells were counted for each sample.

S phase status was monitored by measuring 5-ethynyl-2’-deoxyuridine (EdU) incorporation with the Click-iT™ Plus EdU Alexa Fluor 488 Flow Cytometry Assay Kit (ThermoFisher, C10632) according to the manufacturer’s instructions. 1 μM EdU was added at the time of release to monitor cumulative DNA content between the time of release and the point of fixation. For staining, cells were trypsinised and washed with 3 mL of 1% BSA in PBS, then incubated with 100 μl of Click-iT™ fixative for 15 minutes, pelleted, washed with 3 mL of 1% BSA in PBS and left at 4 °C overnight. Cells were resuspended in 100 μL of 1X Click-iT™ permeabilization and wash reagent and incubated with 500 μl of Click-iT™ Plus reaction cocktail for 30 minutes, before washing once with 3 mL of 1X Click-iT™ permeabilization and wash reagent and resuspended in 500 μL of 1X Click-iT™ permeabilization and wash reagent. FxCycle Violet stain (F10347, Invitrogen) was added to a final concentration of 1 μg/ml. Samples were analysed on a BD LSR II flow cytometer (BD Biosciences) using FACSDiva software (BD Biosciences) and analysed with FlowJo software (BD Biosciences). 10^4^ cells were counted per sample. For quantification the peak of 2N cells was used to gate an assessment of 2N DNA content, as the overlap between S phase and G2/M made it challenging to categorically assign cell cycle status on the basis of DNA content alone. Thus, throughout the manuscript we monitor cell cycle progression as a reduction in 2N DNA content.

### Immunofluorescence

Cells were grown on 13 mm coverslips (No. 1.5, VWR, 631-0150P) in 10 cm dishes (353003, Falcon) using the appropriate growth conditions. For EdU staining, the Click-iT™ EdU Cell Proliferation Kit for Imaging, Alexa Fluor™ 594 dye (Invitrogen, C10339) was used. 1 hour prior to harvesting cells, 10 μM EdU was added to the growth media. Cells were washed in PBS before fixing for 20 minutes in 2% PFA in PBS, before three washes in PBS + 0.1% Tween and storage in PBS + 0.1% Tween at 4 °C overnight. Permeabilisation in PBS + 0.5% Triton-X-100 was followed by 3 washes in PBS + 0.1% Tween. The Click-iT™ reaction for EdU detection according to the manufacturer’s instructions. Cells were washed 3 times in 5% Bovine Serum Albumin (BSA; ThermoFisher Scientific, 11423164) in PBS then incubated with primary antibody to H2AX (Ser139), clone JBW301 (Millipore Cat# 05-636, RRID:AB_309864) at a dilution of 1 in 500 for 1 hour before 3 washes in PBS + 5% BSA and incubation with a 1 in 500 dilution of goat anti-mouse IgG Alexafluor 488 antibody, (Thermo Fisher Scientific Cat# A32723, RRID:AB_2633275), and 2 mg/ml 4’,6-diamidino-2-phenylindole (DAPI, ThermoFisher Scientific, 11916621) for 1 hour. After a further 3 washes in PBS + 0.1% Tween, coverslips were mounted by inversion onto 2 μl of Vectashield Antifade Mounting Medium (Vector Labs H1000). Slides were analysed using an Axioskop2 (Zeiss, Inc) microscope.

### Protein analysis

To collect hTERT-RPE1 protein samples, culture medium was removed, by aspiration, from a 10cm plate, before two washes in PBS and addition of 400 μl TruPAGE LDS sample buffer (Sigma-Aldrich, PCG-3009) containing complete protease inhibitor cocktail (Roche, 11697498001) PhoSTOP (Sigma-Aldrich, 4906837001) and DTT to the plate. Cells were scraped from plate for transfer into a 1.5 ml microfuge tube and snap frozen in liquid nitrogen for storage at −80°C. For THP1, cells were centrifuged at 300 g for 3 minutes and the pellet resuspended in sample buffer as above, frozen in liquid nitrogen and stored at −80°C. Samples were heated at 70°C for 10 minutes before loading on a 10 cm, 10% precast TruPAGE gel (Sigma-Aldrich PCG2009) with TruPAGE SDS running buffer (Sigma Aldrich PCG3001) and transferred to PVDF membrane (BioRad, 1620177) by wet transfer with TruPAGE transfer buffer (Sigma-Aldrich PCG3011). Membranes were blocked in 5% milk and incubated in 2% BSA with primary antibodies to GAPDH (Cell Signaling Technology Cat# 13084, RRID:AB_2713924), Eg5 (Sigma Aldrich Cat# SAB4501650, RRID:AB_10747045), Wee1 (Cell Signaling Technologies Cat# 13084, RRID:AB_2713924) or pHH3 ser10 (custom antibody to peptide ARTKQTARKS*TGGKAPRKQLASK: Eurogentec) overnight at 4°C. After washing, membranes were incubated for 1 hour at room temperature, with the appropriate secondary antibody (Anti-mouse IgG phosphatase conjugated antibody (Sigma-Aldrich Cat# A3688, RRID:AB_258106), or Anti-rabbit IgG phosphatase conjugated antibody (Sigma-Aldrich Cat# A3687, RRID:AB_258103), washed and developed with chromogenic 5-Bromo-4-chloro-3-indolyl phosphate (BCIP, Sigma-Aldrich, B6149).

## Results

### Efficient synchronisation of hTERT-RPE1 cell cycle progression with transient palbociclib treatment

The telomerase immortalised hTERT-RPE1 cell line is widely used for cell cycle and mitotic studies, yet is refractory to double thymidine block induction synchronisation and yields low levels of synchrony following release from serum starvation. We therefore chose this popular line to assess the efficiency of G1 arrest and release by transient exposure to palbociclib as it would be a major benefit to develop synchronisation approaches in this line [56,57].

Cells were grown to 1.5 × 10^4^ cells per cm^2^ in serum supplemented (10%) Dulbecco’s modified eagles medium (DMEM) before release from the substrate with trypsin and plating at a density of 4.4 × 10^3^ cells per cm^2^. 6 hours later palbociclib was added in a range of concentrations from 50 nM to 1 μM to two identical populations. 24 hours later one population was fixed, while the palbociclib containing medium for the other was replaced with pre-warmed medium containing 330 nM nocodazole, before this sample was fixed 24 hours later. DNA content assessment by fluorescence activated cell sorting (FACS) analysis of propidium iodide (PI) stained samples revealed a tight arrest at the 24 hour time point with 2N DNA content at all palbociclib concentrations of 100 nM and higher (Figure 1a,b). Cells readily released into a 4N arrest after being arrested in G1 for 24 h with 100 nM (Figure 1a) and 200 nM palbociclib, however release was less efficient when the arrest was imposed by 500 nM and 1 μM palbociclib (Figure 1b) as the 2N DNA content remained high 24 h after nocodazole addition.

**Figure 1:**
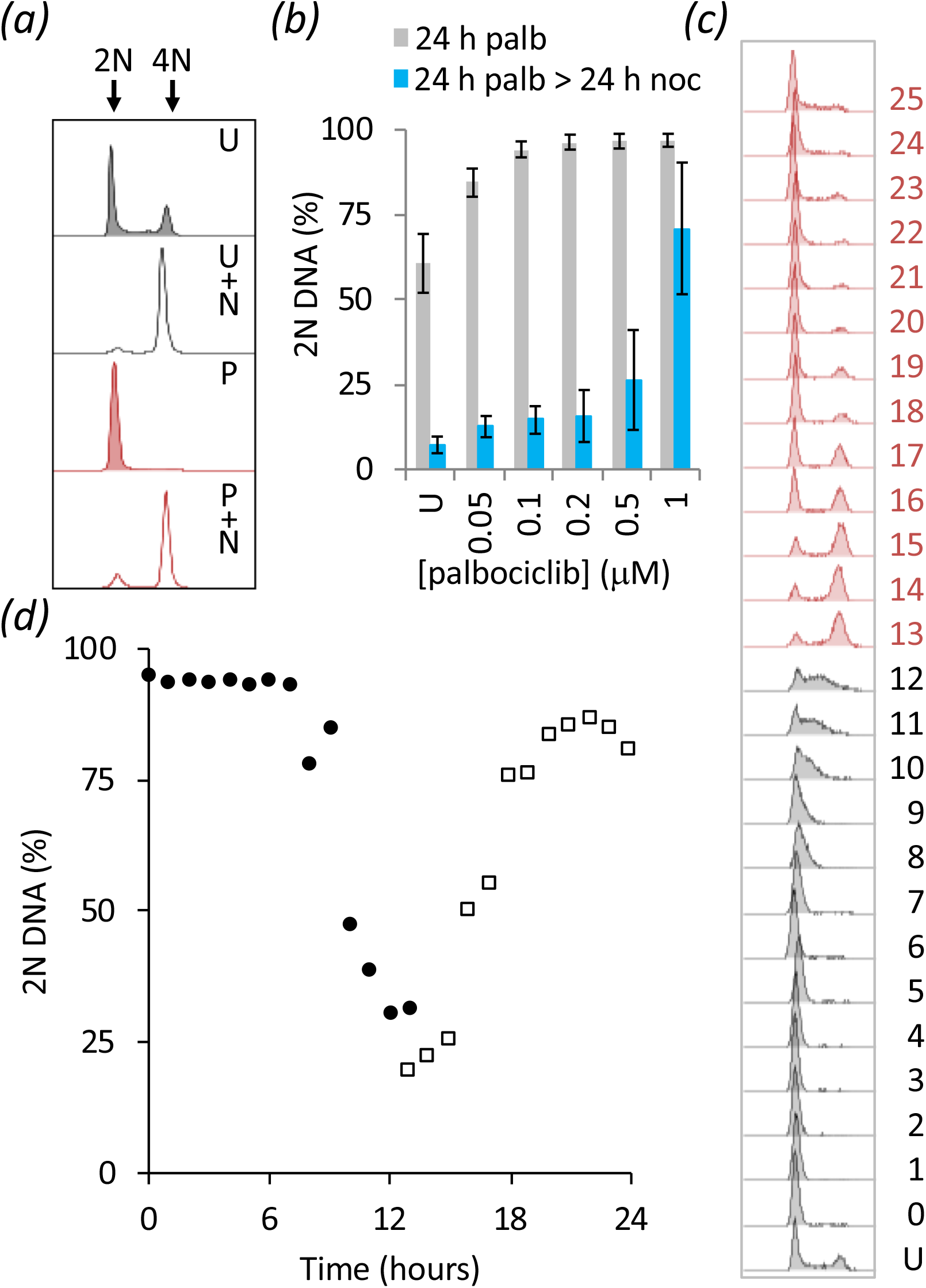
Palbociclib induction synchrony of hTERT-RPE1 cells. (a) hTERT-RPE1 cells were grown to 1.5 × 10^4^ cells cm^2^ in DMEM (+10% serum), trypsinised, and plated at 4.4 × 10^3^ per cm^2^. 6 h later 150 nM palbociclib was added before the culture medium was aspirated and replaced with DMEM + 330 nM nocodazole 24 hours later. Samples were stained for PI FACS analysis at the following time points: just before palbociclib addition (U - untreated), at the switch from palcociclib to nocodazole medium (P) and 24h after this switch to nocodazole (P + N). The strength of the nocodazole induce spindle checkpoint arrest 24 h after the nocodazole was added to an asynchronous population is also shown (U + N). The bimodal peak in the upper panel (U) shows 2N DNA (G1, left) and 4N (G2/M, right) DNA content of an asynchronous, untreated population. (b) Cell populations were treated in the same way as panel (a) with the palbociclib concentration changed to the indicated value. The frequency of 2N cells is plotted at the palbociclib (grey bars) and nocodazole (blue bars) arrest points. Error bars show 1 × SD n = at least 3. (c, d) Population grown as (a) with the exception of release into DMEM without nocodazole and sampling every hour and cells were left for 12 hours after plating, to generate the PI FACS profiles in (c) from which the plots of 2N content shown in (d) were derived. The numbers next to the plots in c indicate time (hours) since release with U indicating an untreated control population. The 13 – 25 h plots (red (c): open squares (d)) were taken simultaneously alongside the 0 - 12 h population (grey (c): filled circles (d)). n = at least 3

Monitoring DNA content at hourly intervals after release from 150 nM palbociclib arrest revealed a synchronous progression from G1 arrest with 2N DNA, through S phase into G2/M phases with 4N DNA content before a return to 2N DNA at 20 hours (Figure 1 c,d). As with all experiments presented herein, 24, or 48, hour periods were covered by sampling parallel populations indicated by black and red in the FACS plots and circles and squares in the graphs. For example, in the 0-13 hour samples of a 24 hour experiment, palbociclib was removed at the start of sampling, while the release had been done 13 hours earlier for samples that were simultaneously collected for the 13-24 hour samples. Consequently, many graphs we present, have 12/13 hour time points from each set.

As the second cycle after release is less likely to be influenced by the physiological challenges of induction synchrony, it is often desirable to monitor this second cycle, rather than the first cycle after release [1]. We therefore extended our assessments to monitor total DNA content over 48 hours after release from 24 hour treatment with 150 nM palbociclib. A notable degree synchrony persisted in the second cycle with 2N cells declining to constitute 46% of the population 30 hours after release (Figure 2a; Supplementary Figure 1). Thus, it will be possible to monitor some trends in biochemical behaviour associated with cell cycle progression in the second cycle after release.

**Figure 2:**
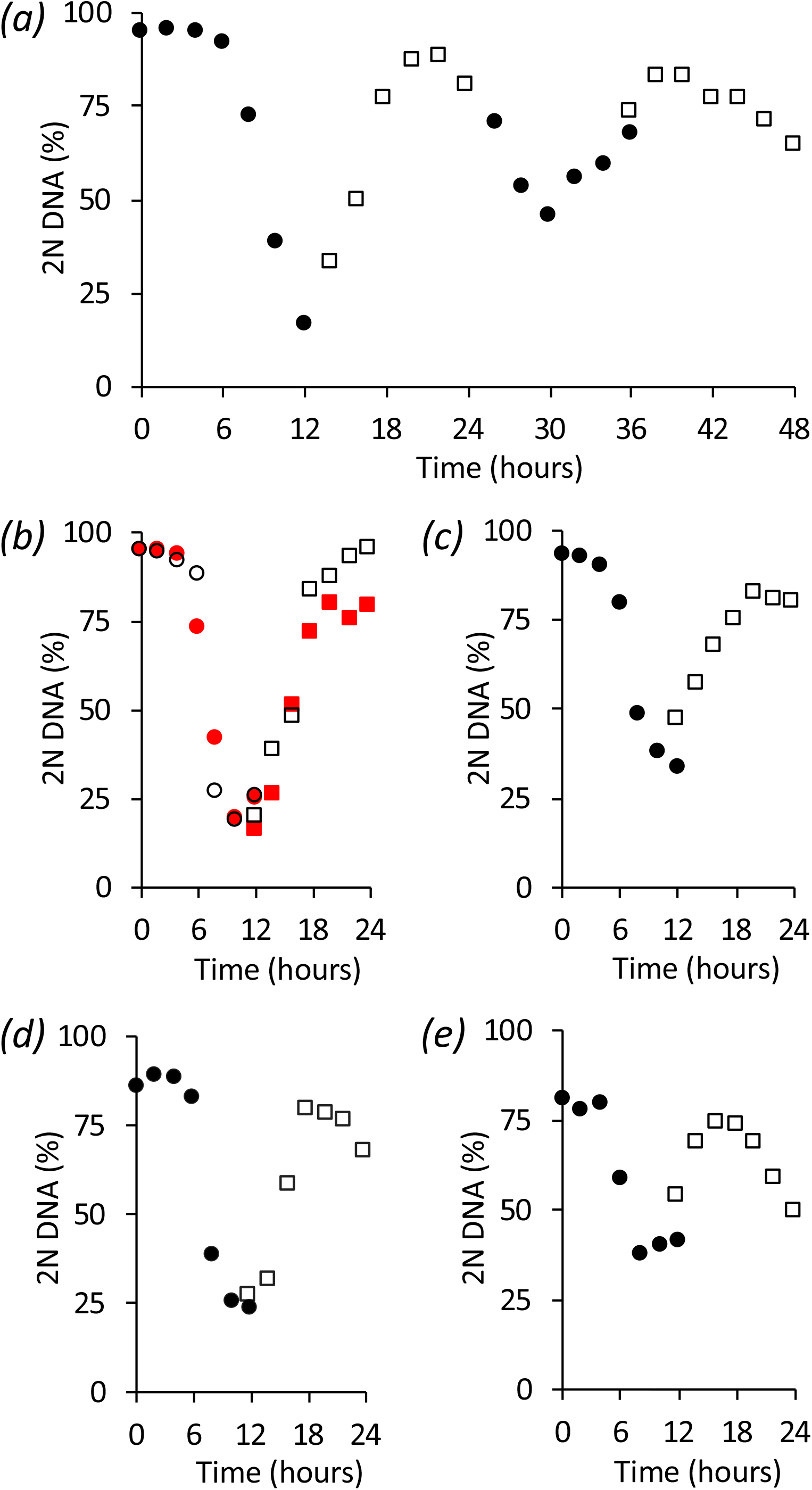
Context and perdurance for palbociclib induction synchronisation. hTERT-RPE1 cells were grown as for Figure 1c with the exception that (a) sampling of 12 hour batches covered a 48 hour release period of the same population of cells. Samples for the 0-12 (filled circles), 14-24 (open squares), 26-36 (filled circles) and 36 to 48 (open squares) were taken in parallel from sub-populations to which the palbociclib had been added at staggered intervals. (b) Two batches of the same population of hTERT-RPE1 cells were followed, in one (open symbols) 150 nM palbociclib was re-added to the population 12h after the initial release from palbociclib, while the other was left to transit the restriction point into the second cycle (red). For (c) the density at which cells from the same population used for (b) were plated was increased 4 fold to 1.76 × 10^4^ cells per cm^2^, while for (d) the expansion of the same starting population used in (b) and (c) lacked one cycle of splitting so that the starting population that was synchronised in (d) had been grown to confluence before plating 12 h before palbociclib addition. (e) Cells were grown in RPMI for two passages before the entire synchronisation outlined in Figure 1c was conducted in RPMI. For (b-e) samples were simultaneously taken from two batches of the same population: palbociclib was added to one at the start of a 12 hour sampling period (circles), while it had been added to the other 12 hours earlier (squares). For the FACS plots from which these 2N DNA contents were derived see Supplementary Figures 1 and 2.

When studying events at the end of the cycle, the progression of the fastest cycling cells in the population into the next cell cycle to initiate a second round of cell cycle events before the events in the first cycle are completely finished in other members of the population, can lead to a merged data dataset in which information from the second cycle is superimposed upon that from the first to obscure finer points of the kinetics of change in the first cycle. We therefore monitored the impact of palbociclib re-addition, after release from the first arrest, to ask whether it would compromise the progression through the observed cycle.

Encouragingly, a second application of Cdk4/6 inhibitor 12 hours after release had no impact upon progression through cycle under study (Figure 2b, compare the red (no re-addition) and black squares (palbociclib added back at 12 h): Supplementary Figure 2a,b). Thus, palbociclib re-addition can be used to insulate observations from consequences of entrance into the next cycle to provide a better insight into the kinetics of cell cycle events.

### Growth conditions are key for optimal synchronisation

The impact of metabolism, growth control, quiescence upon the control of commitment to the cell cycle prompted us to assess the impact of context upon the efficiency of palbociclib induction synchronisation of hTERT-RPE1 cells. There was a notable reduction in the efficiency of synchronisation of hTERT-RPE1 cells when the density of the population seeded onto plastic 6 hours prior to palbociclib addition was increased 4 fold to 1.76 × 10^4^ cells per cm^2^. At this density the proportion of the population with a 2N DNA content only declined to 38% rather than the dip to 16% in the identical population plated at 4.4 × 10^3^ per cm^2^ (Figure 2c: Supplementary Figure 2c). The efficiency of both arrest and release was also compromised when the identical population used in Figures 2b and 2c had been grown to confluence before splitting to generate the populations that were arrested for the synchronisation (declined to only 23% 2N, Figure 2d: Supplementary Figure 2d). Finally, in simultaneous studies of cells from the same initial population that had been passaged twice in Roswell Park Memorial Institute 1640 Medium (RPMI) before synchronisation in this medium, fewer cells arrested cell cycle progression after 24 h in RPMI 1640 (81% vs. 95%) and the proportion of 2N cells only dipped to 37% of the population in RPMI, rather than the decline to 16% in DMEM (Figure 2b, e: Supplementary Figure 2a, e). Thus, culture conditions alter the efficiency of synchronisation and, once initial studies indicate that a line is competent for synchronisation, a variety of conditions should be assessed and care should be taken to ensure that cells remain within active proliferation in the expansion in the lead up to synchronisation.

### Oscillations in established cell cycle markers accompany progression through synchronised cycles

To assess the utility of the approach for biochemical assays, extracts from a palbociclib synchronised culture were probed with antibodies to monitor markers whose levels fluctuate as cells transit cycles synchronised by other means. Anticipated fluctuations in the levels of the kinesin 5 Eg5 and phosphorylation on serine 10 of histone H3 highlight the utility of this approach to monitoring biochemical changes throughout the population (Figure 3).

**Figure 3:**
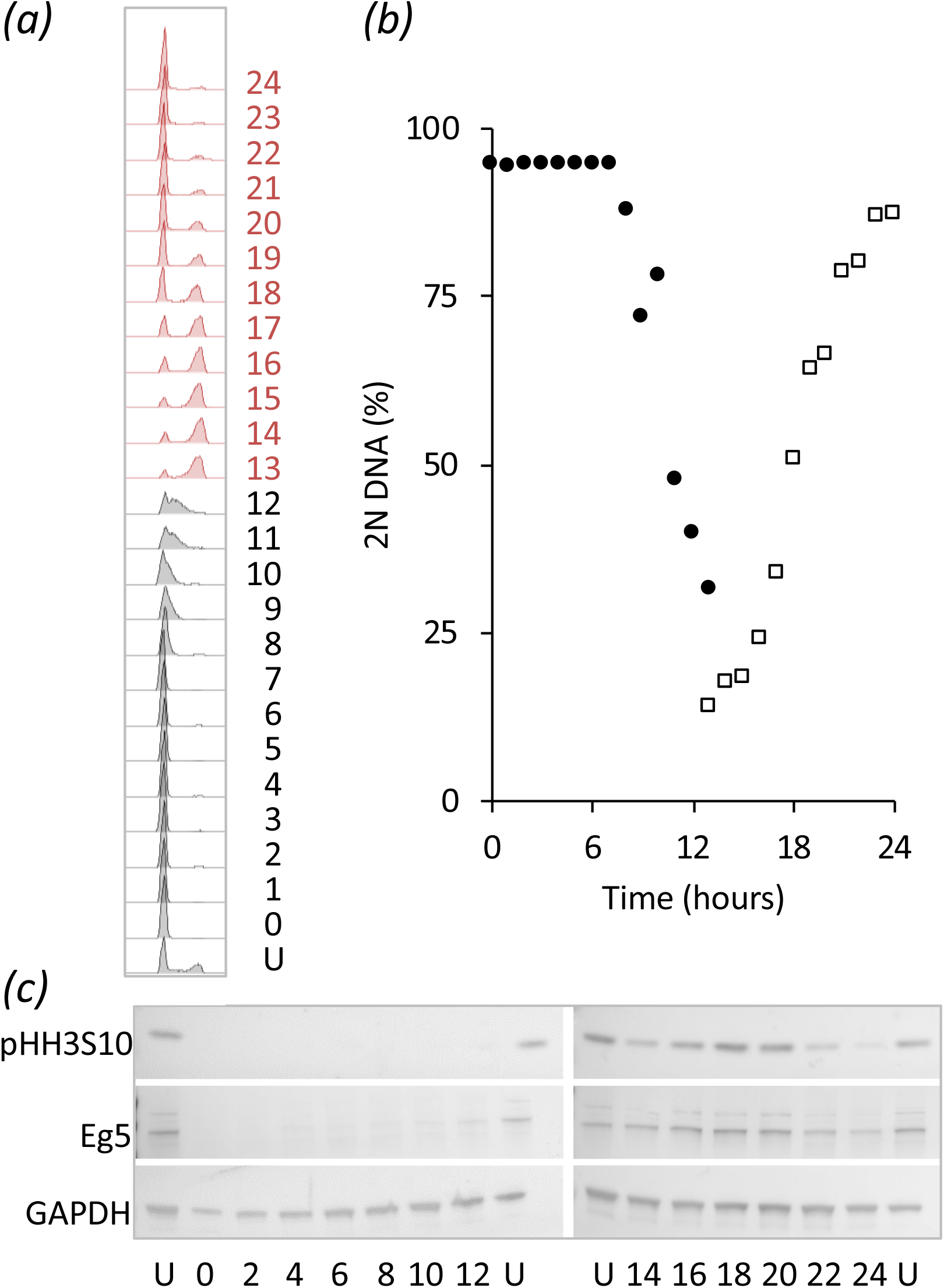
Cdk4/6 induction synchrony can reveal transient cell cycle events. hTERT-RPE1 cells were grown as for Figure 1c and samples taken every hour for PI FACS analysis (a) to gauge the fluctuations in 2N DNA content in the population (b), while sampling to monitor the indicated markers by western blot every 2 hours (c). The numbers next to the plots in (a) indicate hours since release with U indicating an untreated control population. Both the mitotic kinesin 5 motor protein Eg5 and phosphorylation of the serine of histone H3 at position 10 peak as cells return from the 4N state to the 2N state (mitosis and cell division).

### Synchronisation of multiple lines with palbociclib, ribociclib and abemaciclib

We next asked whether palbociclib induction synchronisation would be similarly effective in other lines by assessing the efficiency of arrest and subsequent release into nocodazole arrest of 25 cell lines following exposure to a range of palbociclib concentrations. Although far from exhaustive, these exploratory assessments identified reproducible arrest and release in 5 of the lines tested. We illustrate this efficacy with THP-1, H1299, A549 (Figure 4a-c). The following lines did not show robust arrest/release profiles in our preliminary tests: Hela, U2OS, MCF10A, HCT116, RKO.

**Figure 4:**
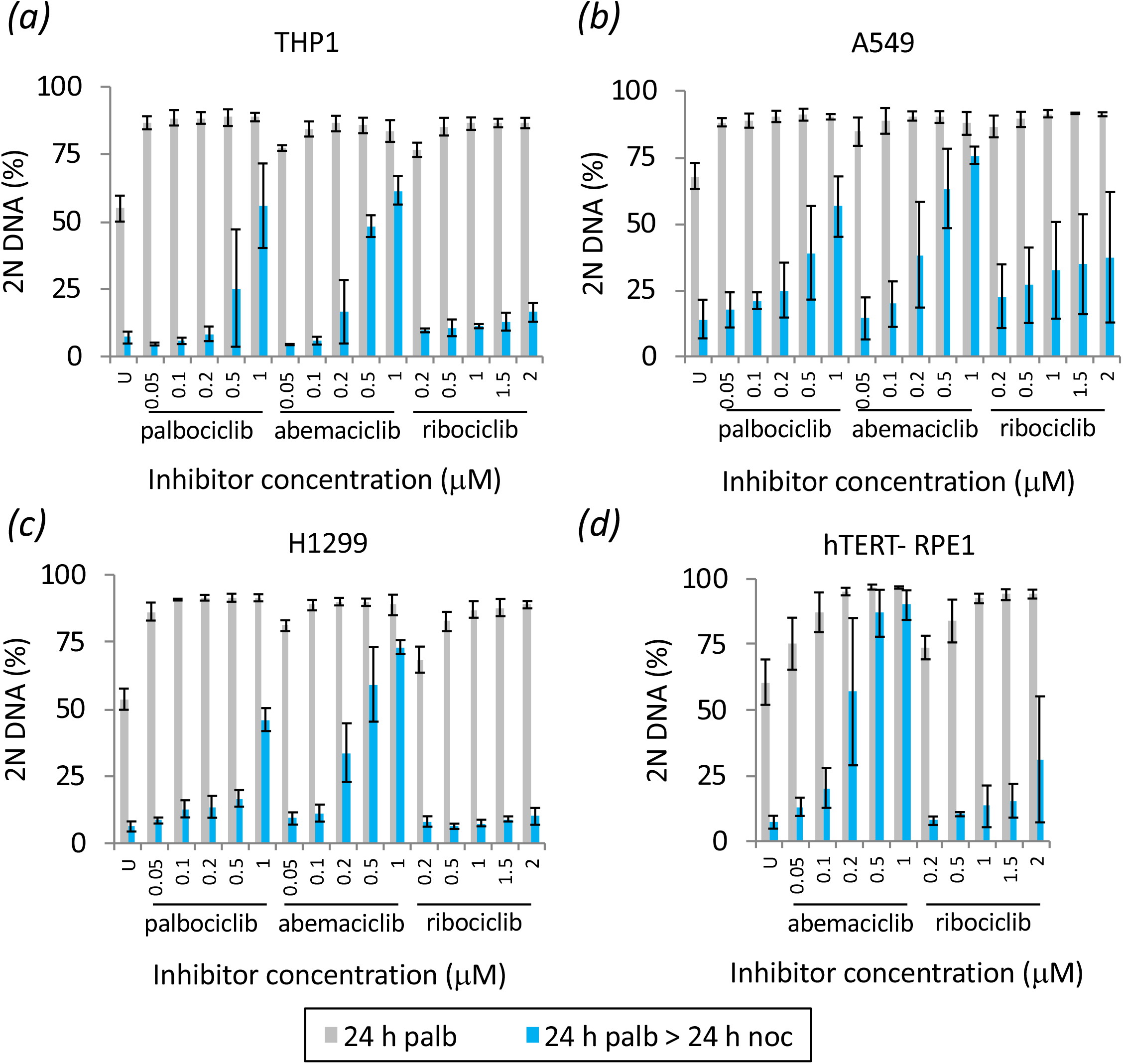
Spectrum of responses of four different lines to three Cdk4/6 inhibitors. The indicated cell lines were all passaged in DMEM and grown as for Figure 1b with the indicated drug concentrations added 6 hours after splitting. Error bars indicate 1 × SD n = at least 3.

As the different Cdk4/6 inhibitors exhibit distinct pharmacological responses [54], we compared induction synchronisation profiles in different lines to gauge the spectrum of responses and whether a reliance upon palbociclib alone as a means to judge competence for CDK4/6 inhibition induction synchrony was a valid approach, or whether other inhibitors may show even greater efficacy. A549, H1299, THP1 and hTERT-RPE1 populations were exposed to a range of palbociclib, ribociclib and abembaciclib concentrations, before the CDK4/6 drug containing medium was swapped for medium containing nocodazole 24 hours into the arrest. These nocodazole containing cohorts were then harvested 24 hours after the swap into nocodazole (Figure 4).

In keeping with the efficacy of palbociclib, these selected lines showed strong Cdk4/6 inhibitor induction synchrony with the two other inhibitors (Figure 4). Palbociclib and ribociclib gave excellent and comparable arrest/release profiles across a broad range of concentrations. In contrast, the window of competence was much narrower for abemaciclib (Figure 4). When the efficiency of G1 arrest is used to identify concentrations of drug that have comparable impact upon restriction point passage, the ability to exit the G1 arrest into the G2 block was markedly lower when the inhibition had been imposed by abemaciclib. For example, for hTERT RPE1 cells, 200 nM of abemaciclib is required to attain the same block in G1 as 100 nm palbociclib, yet after release 57% of the population remained arrested with 2N, rather than the 15 % that persists in the palbociclib treated population (Figure 1b, Figure 4d). This inefficiency is a recurrent theme in all lines (Figure 4a-c). Comparisons between palbociclib and ribociclib reveal subtle distinctions to suggest that, when extensive work is to be performed with a specific line, there would be merit in testing both inhibitors when fine tuning the dose-response profile.

The AML derived line THP-1 is an attractive line for cell cycle studies because cell retrieval via pelleting avoids the challenges of harvesting from adherence to a matrix. We therefore assessed both the robustness of synchrony in this line (Figure 5a, c, d, Supplementary Figure 3) and the behaviour of signature cell cycle markers as a population transited a synchronised culture (Figure 5a,c). As noted for hTERT-RPE1 cells in Figure 3, markers oscillated with their characteristic periodicities (Figure 5b).

**Figure 5:**
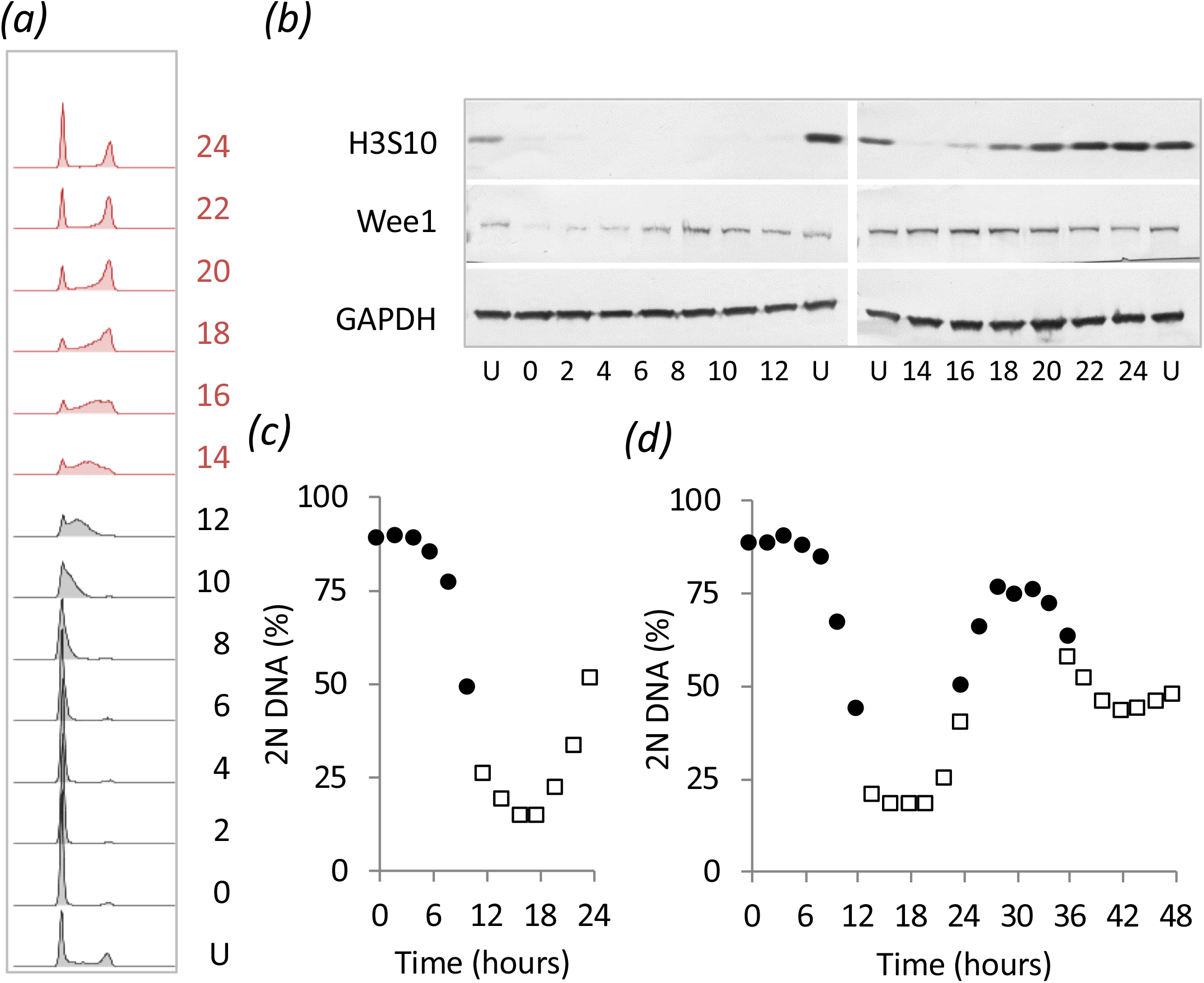
Utility of THP-1 suspension cell line for cell cycle analysis. (a-c) A population of THP1 cells was processed as for Figure 3 with the exception that the isolation of cells for the initial seeding and shift between 100 nM palbociclib medium and drug-free medium was done by mild centrifugation at 300 g for 3 minutes, rather than by trypsinisation, before resuspension in RPMI at a concentration of 1×10^5^ cells/ml. Sampling the population for PI FACS staining (a) to monitor the 2N population (c) was accompanied by isolation of cells for western blotting to detect the proteins indicated in (b). The numbers next to the plots in (a) indicate hours since release with U indicating an untreated control population. (d) Sampling of 12 hour batches covered a 48 hour release period of the same population of cells. Samples for the 0-12 (filled circles), 14-24 (open squares), 24-36 (filled circles) and 36 to 48 (open squares) were taken in parallel from sub-populations to which the palbociclib had been added at staggered intervals. The FACS plots from which the data in (d) are derived are in Supplementary Figure 3.

### Inhibitor cocktails match the efficacy of single agent synchronisation

Off-target inhibition is a concern for interpretation of phenotypes arising from chemical perturbations. With the goal of minimising the off target impacts of each individual inhibitor [54], we asked whether a cocktail of all three inhibitors would be as effective as single agent? We used a mix of one third of the most effective concentration for each individual drug to test the level of cell cycle arrest and release: 33 nM abemaciclib + 33 nM palbociclib + 300 nM ribociclib. Encouragingly, all lines gave robust arrest and efficient release (Figure 6).

**Figure 6:**
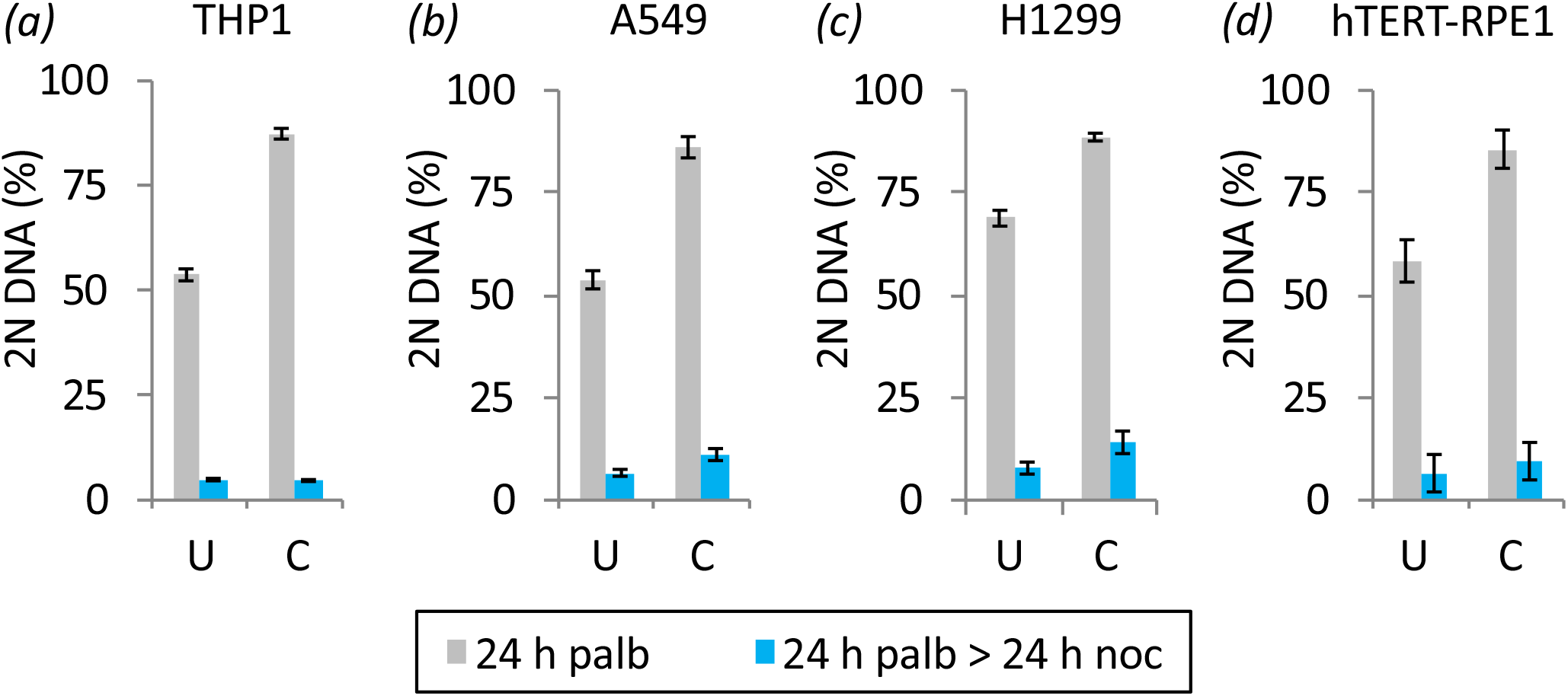
Cocktails support efficient arrest and release. (a-d) Populations of the indicated cell lines were processed as for Figure 1b with the exception that the inhibitor was a cocktail comprising 33 nM abemaciclib + 33 nM palbociclib + 300 nM ribociclib and the media switch for the THP1 was achieved by mild centrifugation at 300 g for 3 minutes. Samples treated with the inhibitor cocktail are indicated by C, while the untreated population by U. Error bars indicate 1 × SD n = at least 3.

### γ-H2AX staining reveals minimal DNA damage in palbociclib induction synchrony

A major limitation of the widely used “double thymidine block” approach is the accumulation of DNA damage during the early S phase arrest [20]. This damage likely arises from collapse, and attempts to repair of the DNA replication forks that stalled because nucleotide provision was compromised [19]. The damage accrued, is likely to account for the abnormal anaphase profiles in the divisions after release from thymidine block [21].

We therefore monitored the accumulation of a marker of DNA double strand breaks, γ - H2AX foci, to assess the level of damage during palbociclib arrest and the ensuing cycle after release. As these foci form naturally during S phase, when replication forks generate double strand breaks, we counterstained cells to identify cells undergoing DNA replication in the hour before sampling by adding 10 μM of the nucleotide 5-ethynyl-2’-deoxyuridine (EdU) one hour before processing each sample. Thus, all EdU positive cells will have been actively replicating DNA in the hour before fixation. This enabled us to distinguish EdU positive cells with γ-H2AX foci, in which we assume the foci are a consequence of DNA replication, from those with no EdU staining in which we assume the foci are indicative of sites of repair of DNA damage. We counted a cell as positive for γ-H2AX foci when immunofluorescence staining revealed more than two foci in a nucleus. To consolidate the insight into the timing of S phase from the assessment of total DNA content throughout the population (Figure 7a, b), we monitored progression through S phase by quantifying the cumulative incorporation of EdU into the DNA after the addition of the lower concentration of 1 μM EdU at the time of release to label of all DNA synthesized after the release. S phase was largely complete by 16 hours after release from 150 nM palbocilib (Figure 7c).

**Figure 7:**
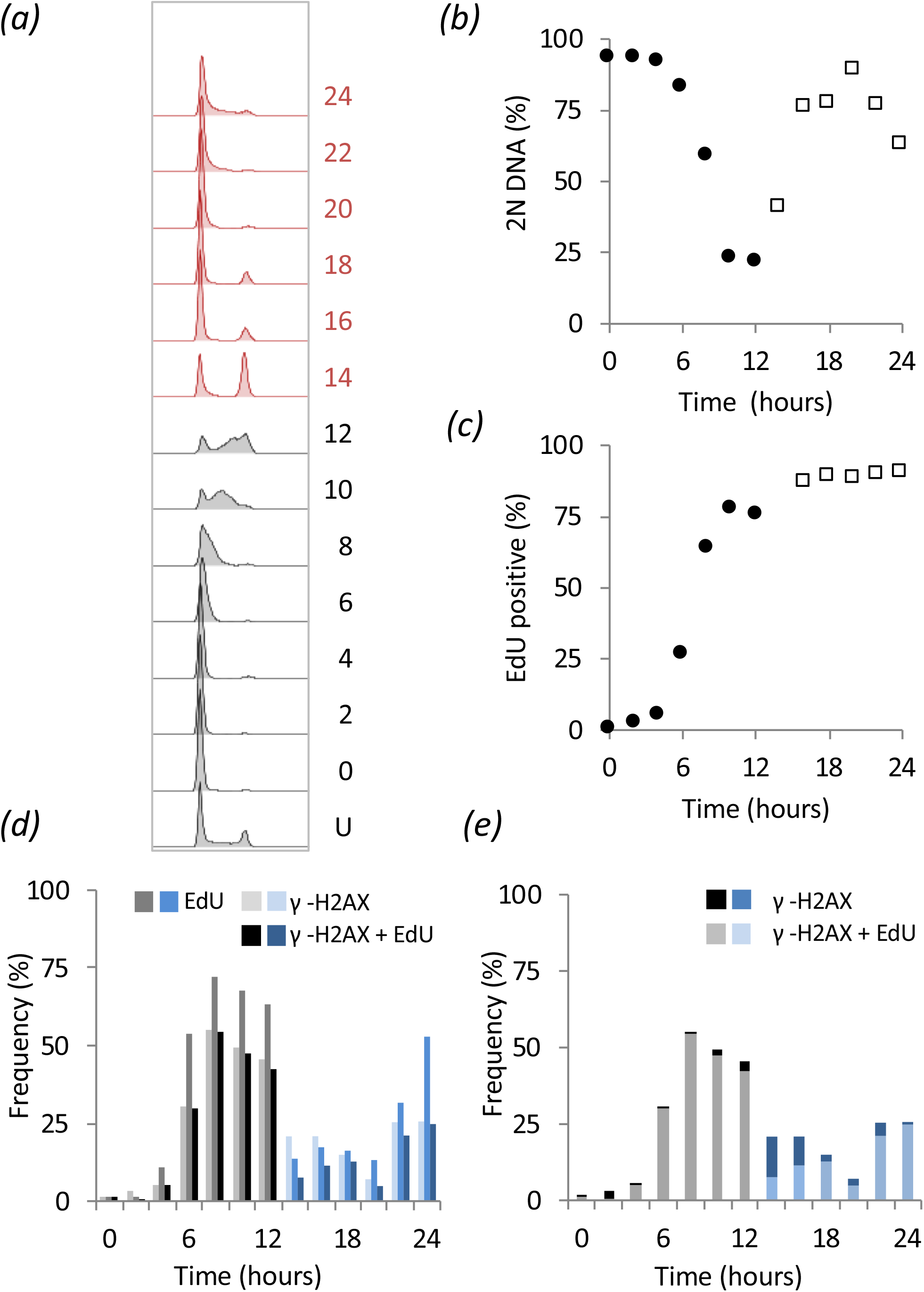
γ-H2AX foci associated with DNA replication in hTERT-RPE1 cells. (a-c) A population of hTERT-RPE1 cells was subjected to arrest release with 150 nM palbociclib treatment as described for Figure 1c and batches stained with PI to measure DNA content (a,b), EdU by addition of 1 μM EdU at the time of release (c), or 10 μM EdU in 1 hour pulses prior to sampling collection alongside antibodies against γ-H2AX (d, e). The numbers next to the plots in (a) indicate hours since release with U indicating an untreated control population. (d) Plots show the frequency of cells with staining of more than two γ-H2AX foci (light fill), EdU (intermediate fill), or those cells positive for both markers (dark fill). (e) γ-H2AX stained cells only: γ-H2AX alone (dark), or both γ-H2AX and EdU (light), at the indicated time points after palbociclib removal. For each time point at least 200 cells were counted to score each characteristic as a proportion of the total counted. The two populations sampled in parallel after staggered palbociclib addition are differentiated by grey and red in (a), closed circles and open squares in (b) and (c), and greys and blues in (d) and (e).

Figure 7 d shows plots from a population of hTERT-RPE1 cells synchronised by palbociclib arrest/release show the frequency of cells incorporating the EdU pulse, those showing two or more γ-H2AX foci and those showing staining with both. For the plots in Figure 7e we just scored cells that contained γ-H2AX foci. The portion of cells in the population with foci, yet no EdU in the darker shading, while those S phase cells staining positive for EdU incorporation are represented by lighter shading (Figure 7e). These plots in Figure 7e show that the vast majority of γ-H2AX positive cells are those replicating. Very few cells in the population had the DNA damage marker, but no EdU staining. Furthermore, cells that have only just initiated S phase may not have incorporated detectable levels of the EdU nucleotide pulse, even though they will have γ-H2AX foci associated with replication forks. Consequently, the assessment by scoring a positive incorporation of the EdU pulse label will give a modest under estimate of S phase cells. Taking this into consideration, and the fact that many cells that incorporated the EdU pulse did not have any γ -H2AX foci (Figure 7d), it would appear that minimal DNA damage accompanies palbociclib induction synchronisation (Figure 7d,e).

As hTERT-RPE1 cells are refractory to synchronisation with thymidine, we used H1299 to directly compare the levels of DNA damage arising during palbociclib induction synchrony and that accumulating at the cell cycle arrest and release following transient addition of 2 mM thymidine to the same starting population (Figure 8, Supplementary Figure 4). Although the degree of synchrony achieved by palbociclib arrest release is not as high in H1299 as in hTERT-RPE1, the results were clear. Consistent with previous reports [20], and in stark contrast to the minimal levels of DNA damage with palbociclib induction synchronisation (Figure 8 a-c), synchronisation by exposure to a single dose of thymidine led to γ-H2AX positive scores for many cells after release from the thymidine arrest point in early S phase (Figure 8 d-f). Importantly, there were foci in many cells that had not incorporated the 1 h EdU pulse label at the time of fixation (the hallmark of actively replicating cells) in cells a long time after the block had been released (Figure 8 e,f). These data suggest that some damage that accumulated at the arrest point persisted throughout the subsequent release, beyond the period of replication. Persistence of damage is consistent with previous reports of chromosomal aberrations during divisions synchronised by thymidine induction synchrony [21].

**Figure 8:**
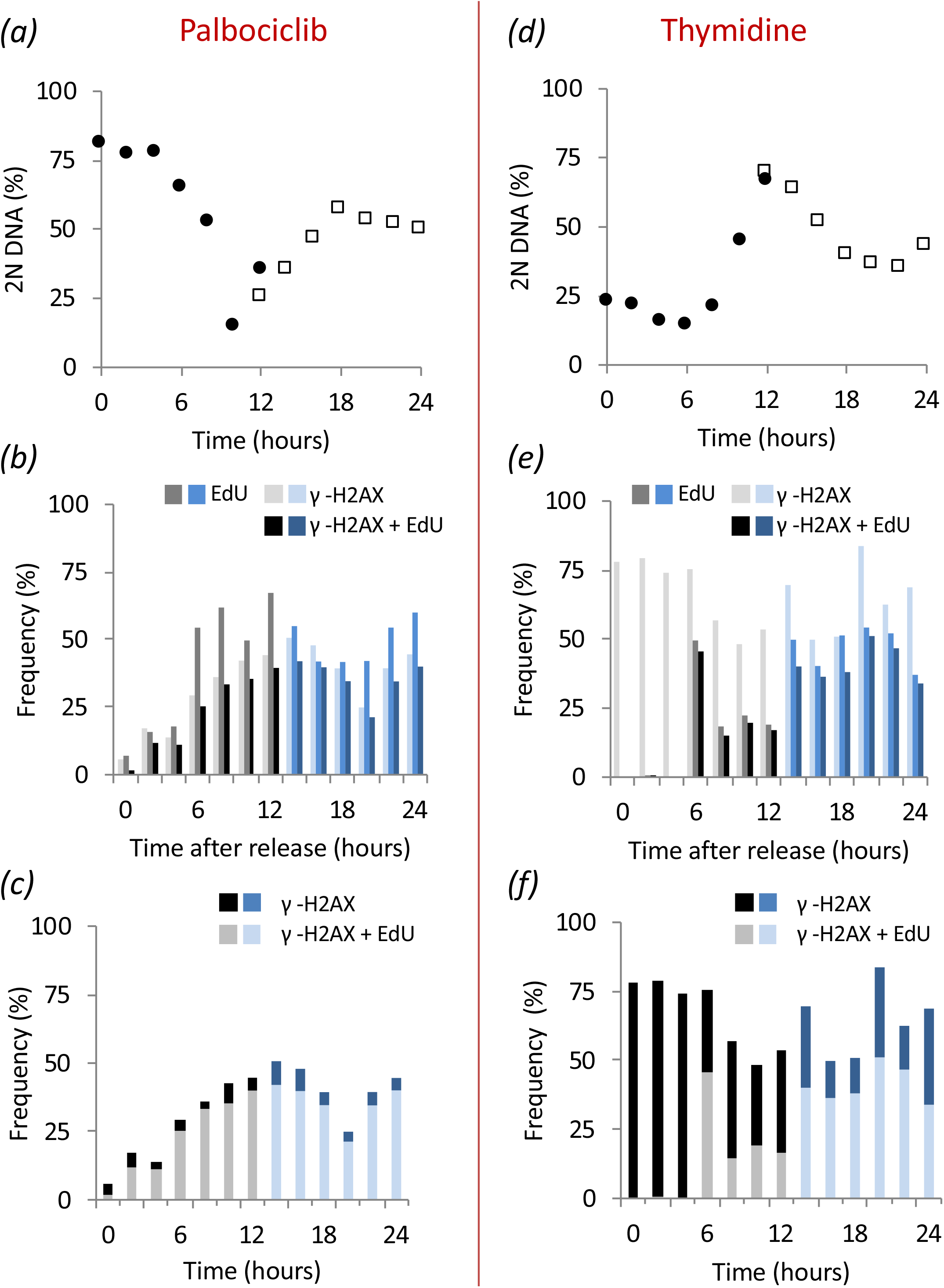
DNA damage at arrest and throughout release in thymidine but not palbociclib synchronised H1299 cells. A population of H1299 cells was subjected to arrest by treatment with 150 nM palbociclib (a-c) or 2 mM thymidine (d-f) and processed as described for Figure 7. The plots in (a) and (d) were derived from the FACs profiles shown in Supplementary Figure 4.

Although we cannot exclude forms of DNA damage that do not generate γ-H2AX foci, palbociclib induction synchronisation of cell cycle progression does not appear to compromise genome integrity to the same degree as thymidine based synchronisation.

### Competence to release maintained over 72 hours

Removal, induction or replacement of a molecule of interest gives great insight into its function. When the manipulation is done in synchronised cultures, it is imperative that the manipulation does not occur in the preceding cycle, otherwise some of the phenotype observed could be an indirect consequence of irrelevant damage accrued in the preceding cycle as cells approach the block point. While impossible to avoid in selection synchronisation, it remains a major challenge in many forms of induction synchronisation that rely upon arrest *within* the cycle. One major appeal of halting cell cycle progression at a point when one cycle is complete and the next is yet to start, is that the impact of any molecular manipulation of the arrested population will generate a phenotype that is a direct consequence of this manipulation upon progression through the ensuing cycle. None of the consequences will be attributable to problems in completing the cycle leading up to the block from which the cells are released.

We therefore compared the competence to return to cycle after Cdk4/6 inhibition for 24, 48 and 72 hours. Pilot experiments established that the ability to hold the arrest, or release after arrest, varied between cell lines and so we selected different palbociclib concentrations to rigorously quantitate the competence to release following protracted arrest (Figure 9). The efficiency of synchronisation was assessed once more by using release into nocodazole to trap cells in the next mitosis as a consequence of activation of the spindle assembly checkpoint [58]. While the ability to maintain the G1 arrest declined to varying degrees in the different lines, with hTERT-RPE1 being the most proficient at maintaining arrest, all lines exited the arrest to reduce the number of 2N cells after 24 hours in nocodazole (Figure 9).

**Figure 9:**
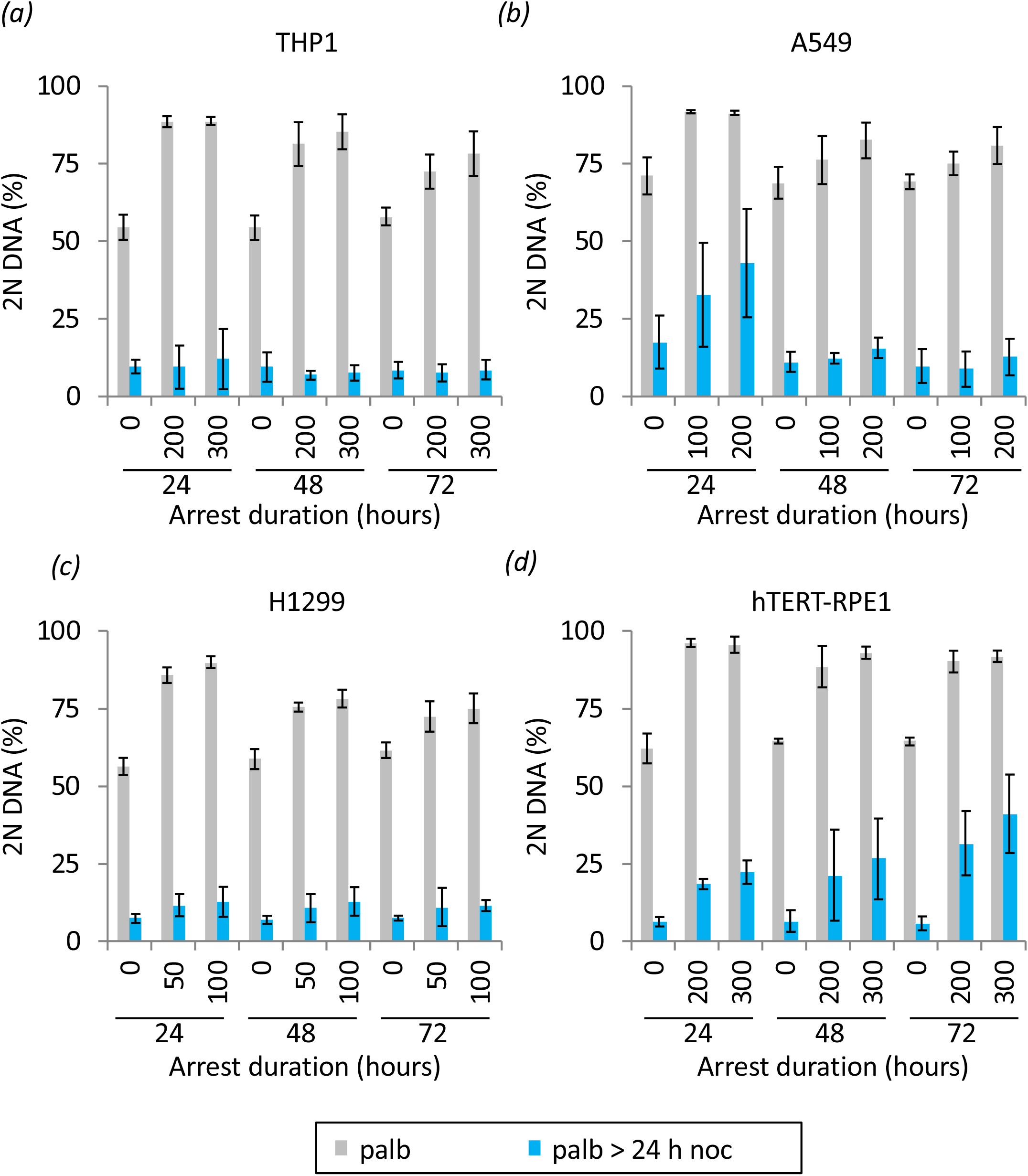
Substantial recovery from extended palbociclib imposed cell cycle arrest. The indicated lines were grown and processed as described for Figure 1b with the exception that cells were maintained in the indicated concentrations of palbociclib for the listed times before wash out into medium containing 330 nM nocodazole. Error bars indicate 1 × SD n = at least 3.

## Discussion

Our examination of the potential of Cdk4/6 inhibition as a novel approach to induction synchrony reveals a highly effective approach to synchronisation of cell cycle progression throughout a population of either adherent, or suspension, human cell lines. We believe that a number of attributes make it a valuable addition to the broad portfolio of cell cycle synchronisation technologies.

The release from palbociclib-imposed cell cycle arrest at the natural decision point for cells, the restriction point, is not associated with changes in growth rates or rates of progression through the cycle into which they are released [25,59]. Rather Cdk4/6-Cyclin D activity appears to set size control [25,59]. Conceptually, this resonates with the original definition of the restriction point, as a rate limiting gateway upon which multiple regulatory systems converge to regulate passage through the gateway into commitment to division [60].

A second appeal of this approach lies in the apparently limited impact upon genome integrity. Although the double thymidine block approach has predominated for over 50 years, the DNA damage that accompanies the arrest persists through the release (figure 8) [20,21]. This cumulative damage limits the utility of this approach for the study in several fields, including DNA replication, DNA repair and chromatin. While assessing genome integrity through the acquisition of γ-H2AX foci is not an exhaustive assessment of damage, our data suggest that Cdk4/6i induction synchrony is not accompanied by the damage and mitotic chromosomal aberrations that arise following release from thymidine [21]. Certainly extensive analyses of mitotic progression following palbociclib induction synchronisation [57] revealed no sign of any of the chromosomal abnormalities that accompany thymidine induction synchronisation (Jon Pines and Mark Jackman personal communication). Cdk4/6i induction synchronisation therefore has the potential to open up a number of previously intractable questions to study in synchronous cultures.

Perhaps the most important appeal of this approach is that cells can remain arrested outside of the cell cycle in full serum over very protracted timescales while retaining the competence to return to a synchronised cycle. As such, this approach will lend itself to protein depletion, induction and molecular replacement while cells are out of cycle for detailed interrogation of the cell cycle modulated attributes of a protein of interest at a level that has been beyond reach to date. Viol and colleagues demonstrate the power of protein induction at the palbociclib arrest point in their study of regulation of centriole biogenesis by the kinase Nek2A [56].

While the benefits of this approach are particularly attractive, as with all approaches to cycle synchronisation, disadvantages will inevitably emerge as these protocols are more widely adopted. Although it seems that cell mass accumulation at the arrest point does not impact upon cell cycle kinetics after release [25], studies in model organisms suggest that there is likely to be adaptation to modified size control at some point after commitment [61]. Recent studies suggest that adaptation is unlikely to occur after the second cycle after release [59]. The approach is also not effective in many lines when the level of control exerted by Cdk4/6 vs Cdk2 is tipped more heavily in favour of Cdk2 control [10,41,47–49,59,62–68].

When a particular line is refractory to Cdk4/6 induction synchronisation, several approaches may switch the line to confer sensitivity to Cdk4/6 inhibition. For lines in which Cdk inhibition alone has little impact, reducing flux through to Cdk4/6 Cyclin D by reducing serum, or inhibiting the downstream MAP kinase pathways can compromise translation to reduce Cyclin D levels and tip the balance to impose cell cycle arrest by palbociclib, or Cdk2 inhibition [10,45,59]. The cell cycle arrest in HCT116 when Trametinib complements palbociclib is a good example of this synergy [69]. The recent revelation that the Cdk4/6 inhibitors are targeting the inactive, rather than active Cdk4 and Cdk6 complexes, suggests another option for refractory lines when resistance arises from greater reliance upon Cdk6, rather than Cdk4 [41]. Cdk4/6 inhibitors impose the arrest because they sequester Cdk4 and Cdk4-CyclinD away from the Hsp90 chaperone system to reduce the number of molecules that can form an active complex with p27 [41,53,70]. The lower affinity of Cdk6 for the Hsp90 chaperone complex enables it to more readily assemble into active trimers that have no affinity for the inhibitors than Cdk4 does. This has prompted the suggestion that the predominance of Cdk6-Cyclin D-p27 trimers depletes p27 from the pools of Cdk2 to elevate Cdk2 activities and confer palbociclib resistance in cancers in which Cdk6 expression is elevated [53,71,72]. Thus, cell lines that rely upon Cdk6 rather than Cdk4 to drive passage through the gateway into the cycle will have reduced sensitivity to Cdk4/6 inhibitors. In such lines, a Cdk6 specific PROTAC may reduce reliance upon Cdk6 to impose sensitivity to Cdk4/6 inhibition and so support synchronisation with the inhibitors [73]. Finally, because many cell lines will bypass the requirement for Cdk4/6 by exploiting Cdk2 to drive cells into cycle, partial inhibition of Cdk2 in these lines [10,74–76] should sensitize cells to Cdk4 inhibition. One challenge with this approach is the accompanying risk that strong Cdk2 inhibition will impact not only upon DNA replication in the preceding cycle, but upon the regulation of mitotic commitment as Cdk2-Cyclin A has been tied directly to regulation of Wee1 and the G2/M transition [10,77–79].

Given the variety of means by which resistance to Cdk4/6 inhibitors arises, unless the genomics indicate a clear resistance to Cdk4/6 inhibition, such as loss of Rb, it is perhaps easiest to empirically test whether a line will be amenable to Cdk4/6i induction synchrony. One of two simple assessments will identify a line as being Cdk4/6i induction synchrony compliant. The arrest/release into nocodazole that we show in Figures 1 and 4 will indicate competence of most lines to synchronise by Cdk4/6i induction synchrony: a line will synchronise if a population accumulates 2N DNA content one doubling time after inhibitor addition, before switching to 4N DNA content a further doubling time after release into nocodazole. However, this assay relies upon the strength of the spindle assembly checkpoint (SAC) [58], that can be so weak in some lines that it fails to impose a long cell cycle arrest with 4N DNA content [80]. The second approach is independent of SAC integrity. Cells are released from palbociclib into 1μM EdU before entrance into the next cycle is blocked by re-addition of palbociclib 12 hours after release. If EdU is incorporated into most genomes throughout a population, that has arrested cell cycle progression with a 2N DNA content in response to the second dose of palbociclib, then the line will be competent for synchronisation.

Our manipulations of cell density and culture history highlight the importance of growth state and culture history when customising the synchronisation protocol to gain maximum synchrony in the chosen line. The plasticity of the G1 control of commitment to the cycle is well documented. A cell’s history alters the configuration of the re-entry switch in both actively cycling cells and those returning from quiescence [45,59,81,82]. Thus, a recent history of contact inhibition is likely to reduce the efficiency of synchronisation, in much the same way it impacts upon the loading of replication factors at origins in hTERT-RPE1 cells [83]. Consequently, care should be exercised to ensure that a population’s path to synchronisation has been freely dividing and as “healthy” as it can be.

While we outline the utility of palbociclib induction synchrony in the study of cell cycle control and execution, pausing the cycle at the restriction point could have considerable benefit in other fields, such as synchronising the formation of the primary cilium by serum depletion from Cdk4/6i arrest, or in studying differentiation in systems, like haematopoesis, where a pause in G1 can be critical for differentiation.

The successful quest to generate drugs to control proliferation in the clinic [51] has generated tools that promise to be of great utility in cell biology and biochemistry laboratories.

## Conclusions

Cdk4/6 induction synchrony is a simple, reproducible, approach that will support biochemical interrogation, induction, depletion and replacement of molecules in any scale of cell line culture. Simple assays can determine competence of this approach to synchronise cell cycle progression in a novel line. A number of approaches can be expected to convert a recalcitrant line into a Cdk4/6i synchronisation compliant line.

## Author Contributions

IH conceived the study and designed the experiments with ET who executed every experiment before IH wrote the manuscript with support from ET.

## Acknowledgements

We thank: Andrew Hudson and Cynthia Li for assistance with sample collection, Jeff Barry and Toni Banyard from the CRUK Manchester Institute Flow Cytometry core facility for support with FACS analysis, Tony Carr and Jon Pines for stimulating discussions and Stephen Taylor, Anthony Tighe, Christine Schmidt for guidance on γ-H2AX staining. We thank Mark Jackman and Jon Pines for communicating their data on mitotic progression following release from palbociclib arrest. IH and ET are supported by CRUK grant (A27336 and A24458).

## Supplementary Material

**Supplementary Figure 1: PI FACS profiles of synchronised populations in Figure 2a** The plots of PI FACS analysis of the populations shown in Figure 2a. The parallel samples indicated by alternating filled circles and open squares in Figure 2a are distinguished by alternating grey and red shading. Numbers indicate hours since release while U = untreated.

**Supplementary Figure 2: PI FACS profiles of synchronised populations in Figure 2b-e** The plots of PI FACS analysis of the populations shown in the respective panels of Figure 2b-e. The parallel samples indicated by filled circles and open squares in Figure 2 are distinguished by the grey and red shading. Numbers indicate hours since release while U = untreated.

**Supplementary Figure 3: PI FACS profiles of synchronised populations in Figure 5d** The plots of PI FACS analysis of the populations shown in Figure 5d. The parallel samples indicated by alternating filled circles and open squares in Figure 5d are distinguished by alternating grey and red shading. Numbers indicate hours since release while U = untreated.

**Supplementary Figure 4: PI FACS profiles of synchronised populations in Figure 8** The plots of PI FACS analysis of the populations shown in the respective panels of Figure 8. The parallel samples indicated by filled circles and open squares in Figure 8 are distinguished by the grey and red shading. Numbers indicate hours since release while U = untreated.

