## Supplementary Figures for "Release from cell cycle arrest with Cdk4/6 inhibitors generates highly synchronised cell cycle progression in human cell culture"

Trotter and Hagan Supplementary Figure 1

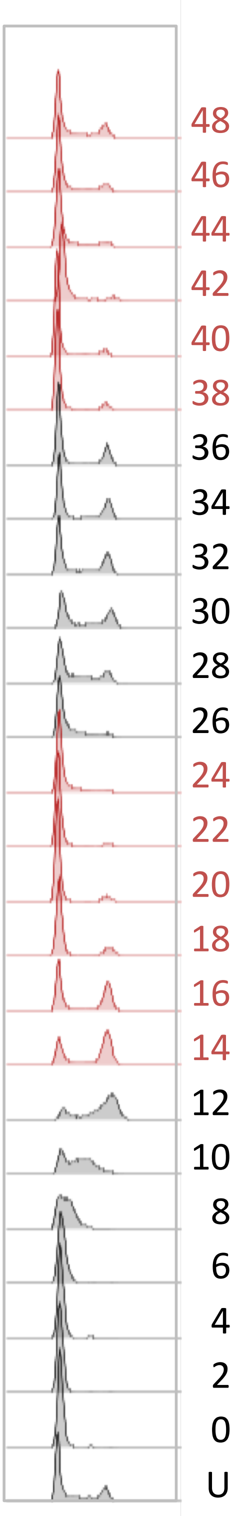


48 hour hTERT - RPE1

Trotter and Hagan Supplementary Figure 2

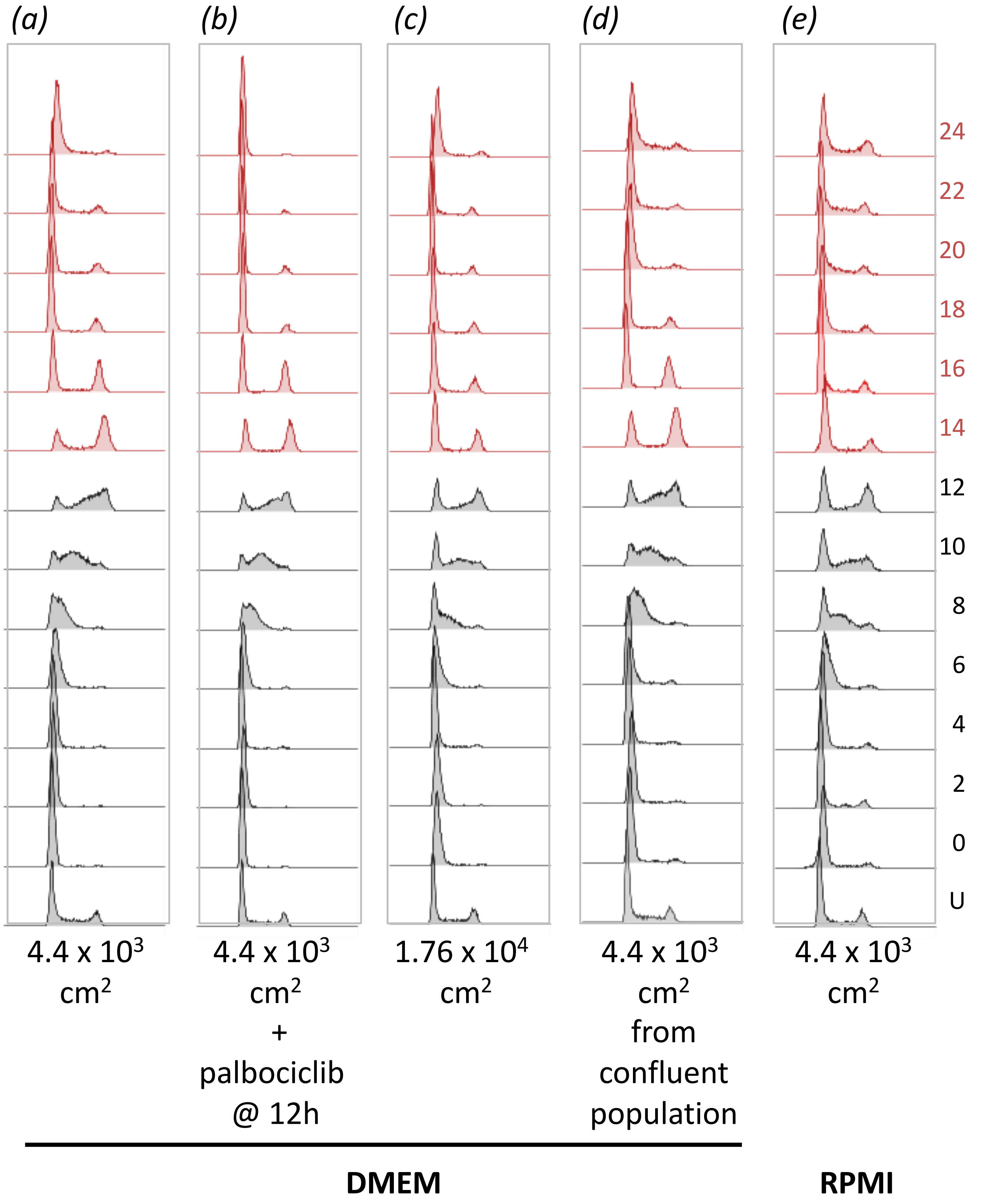

### Trotter and Hagan Supplementary Figure 3

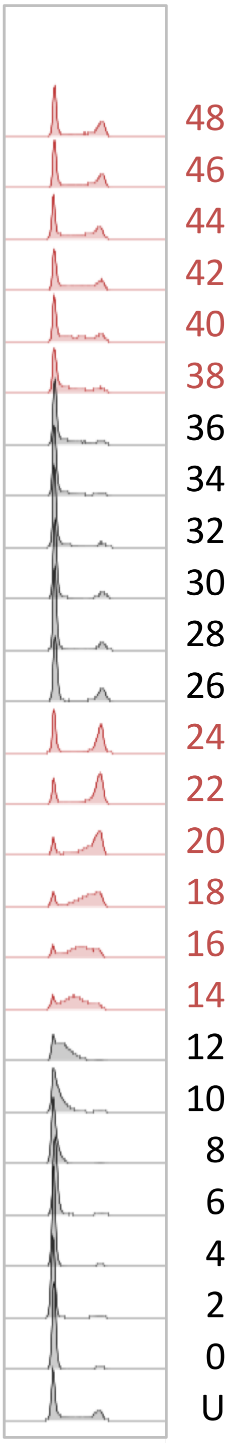


48 hour THP1

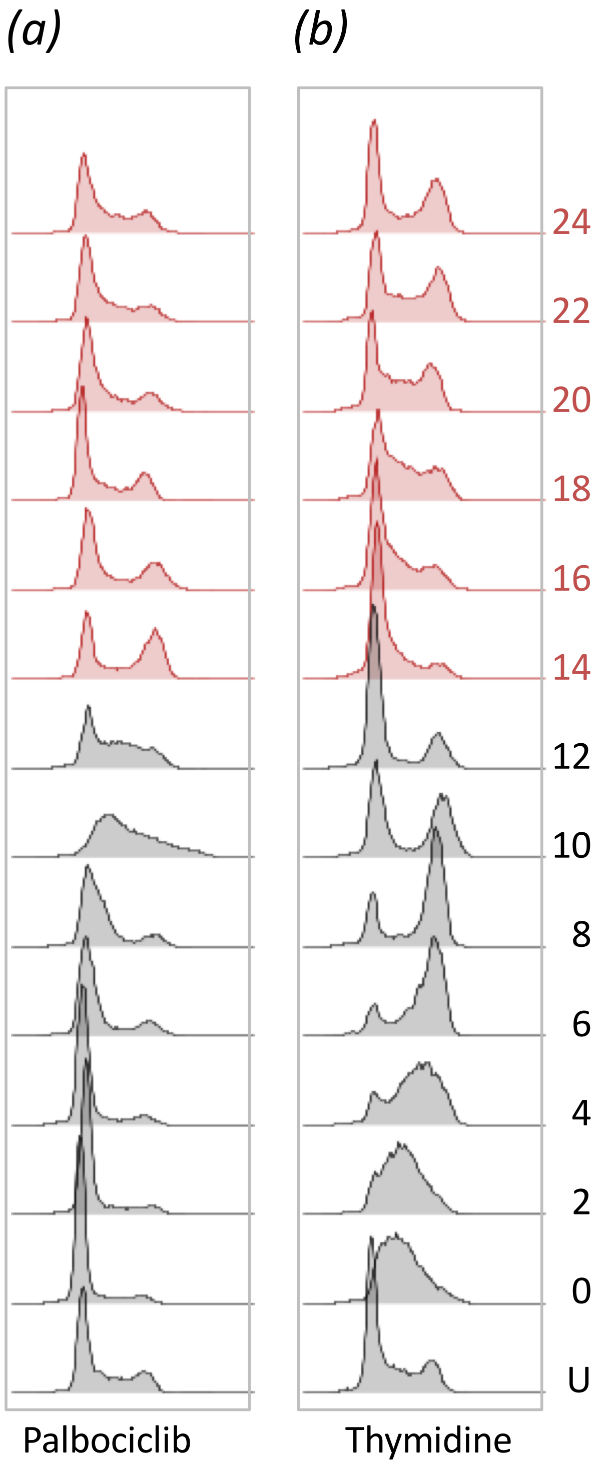
(a)
(b)
24
22
20
18
16
14
12
10
8
6
4
2
0
U
Palbociclib
Thymidine
